# PIN-LIKES coordinate brassinosteroid signalling with nuclear auxin input in *Arabidopsis thaliana*

**DOI:** 10.1101/646489

**Authors:** Lin Sun, Elena Feraru, Mugurel I. Feraru, Krzysztof Wabnik, Jürgen Kleine-Vehn

**Author notes:** Correspondence should be addressed to J.K.-V.

## Abstract

Auxin and brassinosteroids (BR) are crucial growth regulators and display overlapping functions during plant development. Here, we reveal an alternative phytohormone crosstalk mechanism, revealing that brassinosteroid signaling controls nuclear abundance of auxin. We performed a forward genetic screen for *imperial pils* (*imp*) mutants that enhance the overexpression phenotypes of PIN-LIKES (PILS) putative intracellular auxin transport facilitator. Here we report that the *imp1* mutant is defective in the brassinosteroid-receptor BRI1. Our data reveals that BR signaling transcriptionally and posttranslationally represses accumulation of PILS proteins at the endoplasmic reticulum, thereby increasing nuclear abundance and signaling of auxin. We demonstrate that this alternative phytohormonal crosstalk mechanism integrates BR signaling into auxin-dependent organ growth rates and likely has widespread importance for plant development.

## Introduction

The phytohormone auxin is a key regulator of plant development. Indole-3-acetic acid (IAA), the most abundant endogenous auxin, is perceived by the nuclear F-Box protein TRANSPORT INHIBITOR RESPONSE 1 (TIR1) and its close homologs (Dharmasiri et al., 2005a; Kepinski and Leyser, 2005). Auxin facilitates the binding of TIR1 to its co-receptors of the AUXIN/INDOLE ACETIC ACID (Aux/IAA) family, which initiates the proteasome-dependent degradation of the latter. Subsequently, the AUXIN RESPONSE FACTORs (ARFs) are released from the inhibitory heterodimerization with Aux/IAAs and trigger transcriptional responses (Weijers and Friml, 2009). The TIR1 pathway is also involved in rapid, non-genomic responses (Fendrych et al., 2018), but the underlying mechanism remains to be elucidated.

Most IAA is synthesized in a two-step biosynthetic route, providing auxin in various tissues (Mashiguchi et al., 2011; Phillips et al., 2011; Won et al., 2011). Additionally, plants evolved several mechanisms that are thought to, either transiently (auxin conjugation and conversion) or irreversibly (auxin oxidation and conjugation to certain moieties), modify auxin molecules (Ostin et al., 1998; Staswick et al., 2005; Kai et al., 2007; Peer et al., 2013; Pencik et al., 2013). These molecular modifications of IAA ultimately abolish its binding to TIR1, thereby directly impacting the nuclear auxin signaling rates (Sauer et al., 2013).

Besides local auxin metabolism, intercellular auxin transport is crucial to define auxin signaling gradients and maxima within plant tissues (Benkova et al., 2003). The canonical, plasma membrane-localized PIN-FORMED (PIN) auxin efflux facilitators mainly determine the directionality of intercellular auxin transport and, hence, have outstanding developmental importance (Wisniewska et al., 2006). Intriguingly, non-canonical PIN auxin facilitators, such as PIN5 and PIN8, are at least partially retained in the endoplasmic reticulum (ER) and indirectly modulate auxin signaling, presumably through an ER auxin sequestration mechanism (Mravec et al., 2009; Dal Bosco et al., 2012; Ding et al., 2012; Sawchuck et al., 2013).

In an *in-silico* screen, we identified the PIN-LIKES (PILS) protein family of auxin transport facilitators, which resembles the predicted topology of PIN proteins (Barbez et al., 2012). Despite some structural similarities, the evolution of PIN and PILS proteins is nevertheless distinct within the plant lineage (Barbez et al., 2012; Feraru et al., 2012). At the subcellular level, PILS putative auxin facilitators control the intracellular auxin accumulation at the ER and restrict nuclear availability and signaling of auxin (Barbez et al., 2012; Feraru et al., 2012; Beziat et al., 2017; Feraru et al., 2019). Thereby, PILS proteins determine the cellular sensitivity to auxin and contribute to various growth processes during plant development (Barbez et al., 2012; Beziat et al., 2017; Feraru et al., 2019).

*PILS* transcription is highly sensitive to environmental fluctuations, such as light and temperature, integrating external signals to modulate auxin-dependent growth rates (Beziat et al., 2017; Feraru et al., 2019). Using a forward genetic screen, we reveal here that *PILS* genes also function as important integrators of endogenous cues, such as brassinosteroid (BR) hormone signaling. Our work illustrates that BR signaling restricts *PILS* transcription and protein abundance and, thereby, increases nuclear abundance and signaling of auxin. We conclude that this alternative phytohormonal crosstalk mechanism, integrating BR signaling with auxin-dependent organ growth rates.

## Results

### Impaired BR perception enhances PILS5 overexpression phenotypes

To assess how intracellular PILS auxin transport facilitators regulate plant development, we performed an unbiased, forward genetic screen. We used an ethyl methanesulfonate (EMS)-mutagenized population of a constitutively expressing PILS5 line (*35S::PILS5-GFP*/*PILS5*^*OE*^) and screened for mutants that either enhance or suppress PILS5-related dark-grown hypocotyl phenotypes (Figure 1A). *PILS5*^*OE*^ seedlings show shorter, partially agravitropic hypocotyls, and premature apical hook opening in the dark (Barbez et al., 2012; Beziat et al., 2017; Figure 1B, 1C). From more than 3,000 M1 families, we identified 8 *imperial pils* (*imp*) mutants that markedly enhanced the PILS5-related dark grown hypocotyl phenotypes. Here, we describe the *imp1* mutation, which did not only severely impact on PILS5-dependent hypocotyl growth in the dark (Figure 1B, 1C), but also augmented defects in main root expansion in light-grown seedlings (Figure 1D, 1E), suggesting a broad impact on PILS5-reliant traits.

**Figure 1.**
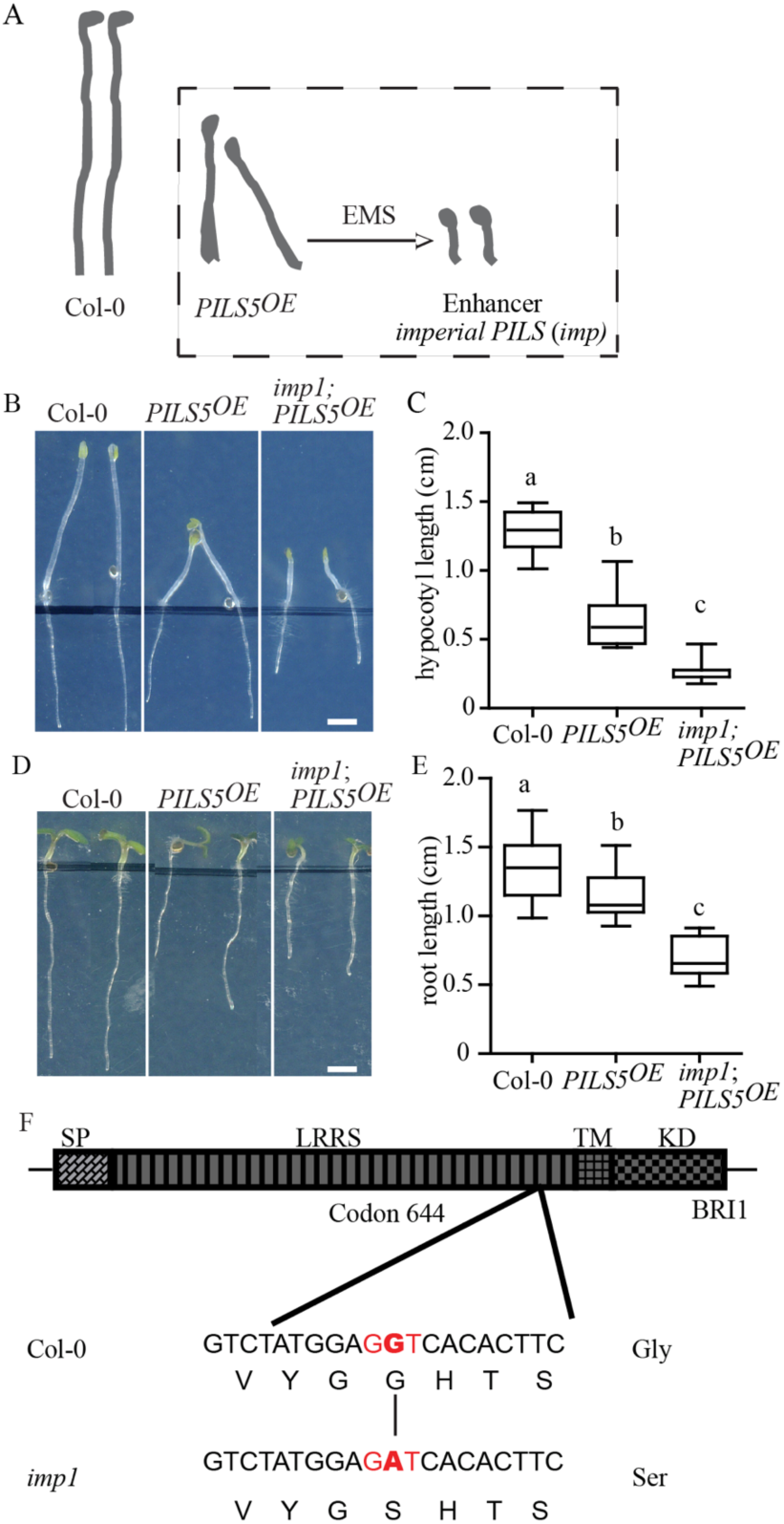
*imp1* mutation enhances PILS5 overexpression phenotypes. A, schematic diagram depicts the “EMS enhancer screen” for identification of genetic modulators of PILS5-related traits. B-E, images and quantifications of 4-d-old dark-grown and 6-d-old light-grown seedlings of *Col-0, PILS5*^*OE*^ and *imp1* mutant grown on ½ MS Scale bar, 30 mm. (n > 25). Letters indicate values with statistically significant differences (P < 0.01, one-way ANOVA). F, sketch of *imp1* mutation in the *BRI1* locus. The diagram shows the full-length BRI1 protein with a defined signal peptide (SP), leucine-rich repeat (LRR), transmembrane (TM), and kinase (KD) domain. The change of G to A in *imp1* results in the conversion of glycine (G) to serine(S) at amino-acid residue 644 in the LRR domain of BRI1.

To identify the underlying mutation, we used a combination of classical mapping and next generation sequencing (NGS). During rough mapping, the *imp1* mutation associated within a region of chromosome 4 (18.096Mb-18.570Mb), where NGS identified a single mutation (Adenine to Guanine) that resulted in an amino acid change (G644 to S) in the brassinosteroid receptor *Brassinosteroid Insensitive 1* (*BRI1*) (Figure 1F). The identified mutation is reminiscent to the previously isolated allele *bri1-6* or *bri1-119*, which altered the same site (G644 to D) (Noguchi et al., 1999; Friedrichsen et al., 2000). In agreement, *imp1;PILS5*^*OE*^ rosettes largely resembled the *bri1* mutant phenotype (Figure 2A), proposing that the *imp1;PILS5*^*OE*^ mutant is impaired in BR signaling.

**Figure 2.**
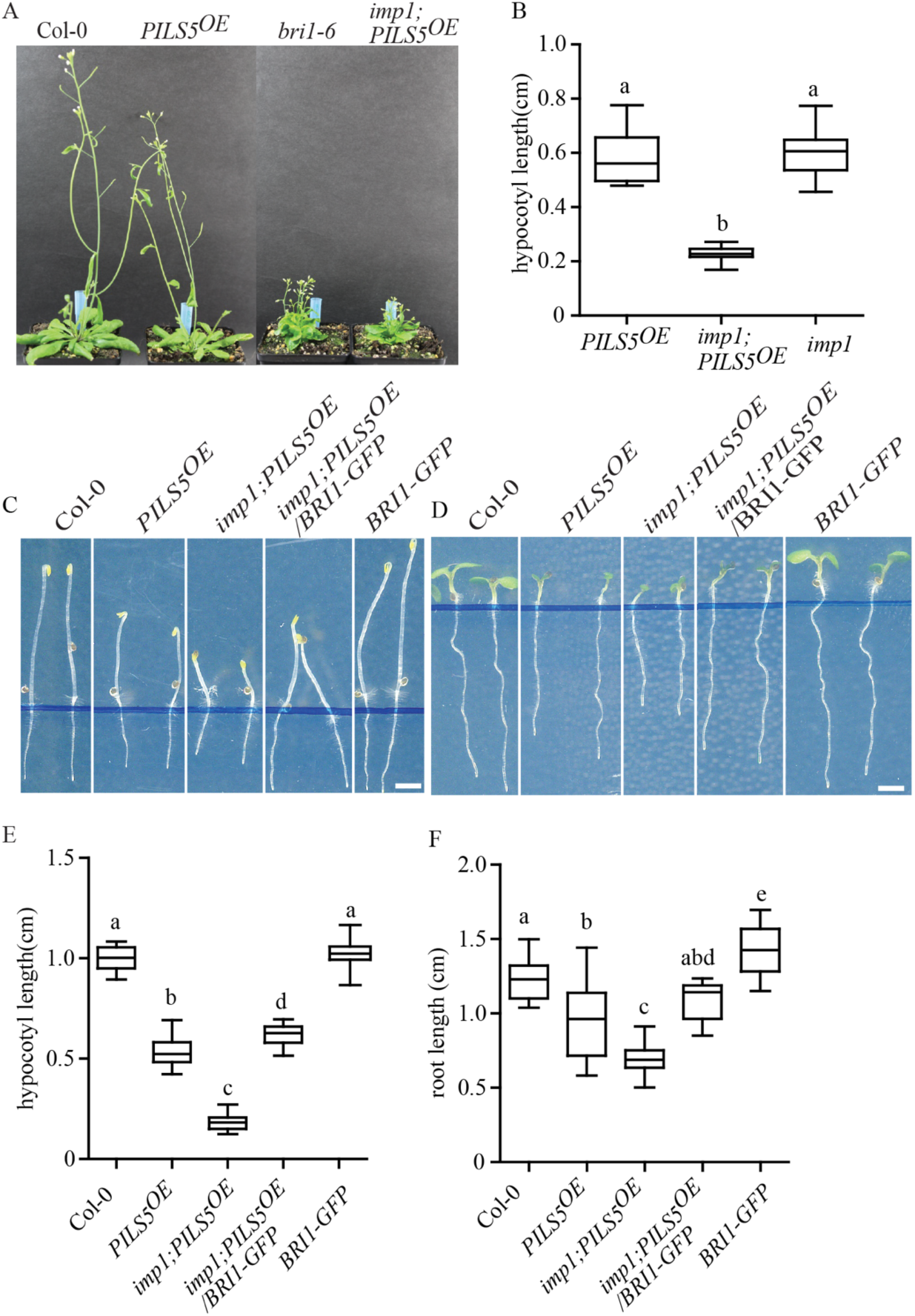
Impaired BR perception impacts on PILS5-related phenotypes. A, 6-w-old plants of wild type, *PILS5*^*OE*^, *bri1-6*, and *imp1;PILS5*^*OE*^ under standard growth conditions. B, 5-d-old dark-grown hypocotyls quantifications of *PILS5*^*OE*^, *imp1*, and *imp1;PILS5*^*OE*^ mutants (n > 25). C-F, images and quantifications of 5-d-old dark-grown (C, E) and 6-d-old light-grown (D, F) seedlings of wild type and indicated mutant lines. Scale bar, 30 mm. (n > 25). w: weeks. Letters indicate values with statistically significant differences (P < 0.01, one-way ANOVA).

To phenotype the *imp1* mutant independently of *PILS5*^*OE*^, we outcrossed the *imp1* mutation to *Col-0* wild type twice and revealed that the *imp1* mutant showed a similar reduction in dark-grown hypocotyl length as *PILS5*^*OE*^, confirming a strong additive effect in *imp1;PILS5*^*OE*^ mutant combination (Figure 2B). Next, we tested the BR sensitivity of *imp1* mutant seedlings. Similar to *bri1-6*, the dark-grown hypocotyls as well as the light-grown roots of *imp1* mutant were strongly resistant to application of 24-Epibrassinolide (BL) (Supplemental Figure 1A-1D). These findings confirm that *imp1* mutant seedlings are impaired in BR signaling.

To further test whether the absence of *BRI1* enhances PILS5-related phenotypes, we expressed *pBRI1::BRI1-GFP* in the *imp1;PILS5*^*OE*^ mutant background. BRI1-GFP expression indeed complemented the *imp1;PILS5*^*OE*^ mutant, resembling *PILS5*^*OE*^ phenotypes (Figure 2C-2F). This data suggests that the *bri1*^*imp*^ mutation is responsible for the enhanced *PILS5*^*OE*^-related phenotypes. Additionally, overexpression of *PILS5* in the *bri1-6* mutant background largely phenocopied the *imp1;PILS5*^*OE*^ mutant seedlings (Supplemental Figure 1E-1H). This set of data suggests that BR perception indeed impacts on PILS5-related traits.

### BR signaling modulates *PILS* gene expression and *PILS* protein turnover

We next investigated if BR signaling modulates *PILS* gene activity, because in silico analysis revealed potential binding sites for BR-dependent transcription factors BZR1 and BZR2 in the promoters of *PILS2, PILS3* and *PILS5* (Supplemental Figure 2). Moreover, based on chromatin immunoprecipitation (ChIP)-sequencing, *PILS2* as well as *PILS5* are direct targets of BZR1 (He et al., 2005; Chaiwanon and Wang, 2015). Exogenous application of BL repressed the transcriptional reporters *PILS2, PILS3*, and *PILS5* fused to green fluorescent protein (GFP) and β-glucuronidase (GUS) (Figure 3A-3F). Furthermore, *pPILS5::GFP-GUS* was reduced and enhanced in roots of *BRI1* overexpressing lines and in roots of *bri1* mutant alleles, such as *bri1-5* (Noguchi et al., 1999), *bri1-6* as well as *imp1*, respectively (Figure3G, 3H; Supplemental Figure 3A-3D). Moreover, we also detected reduced *pPILS5::GFP-GUS* activity in roots of constitutively active *bzr1d* (Figure 3I, 3J), which suggests that transcription factor BZR1 negatively regulate *PILS5* expression. Based on these findings, we conclude that BR signaling limits the transcription of *PILS* genes.

**Figure 3.**
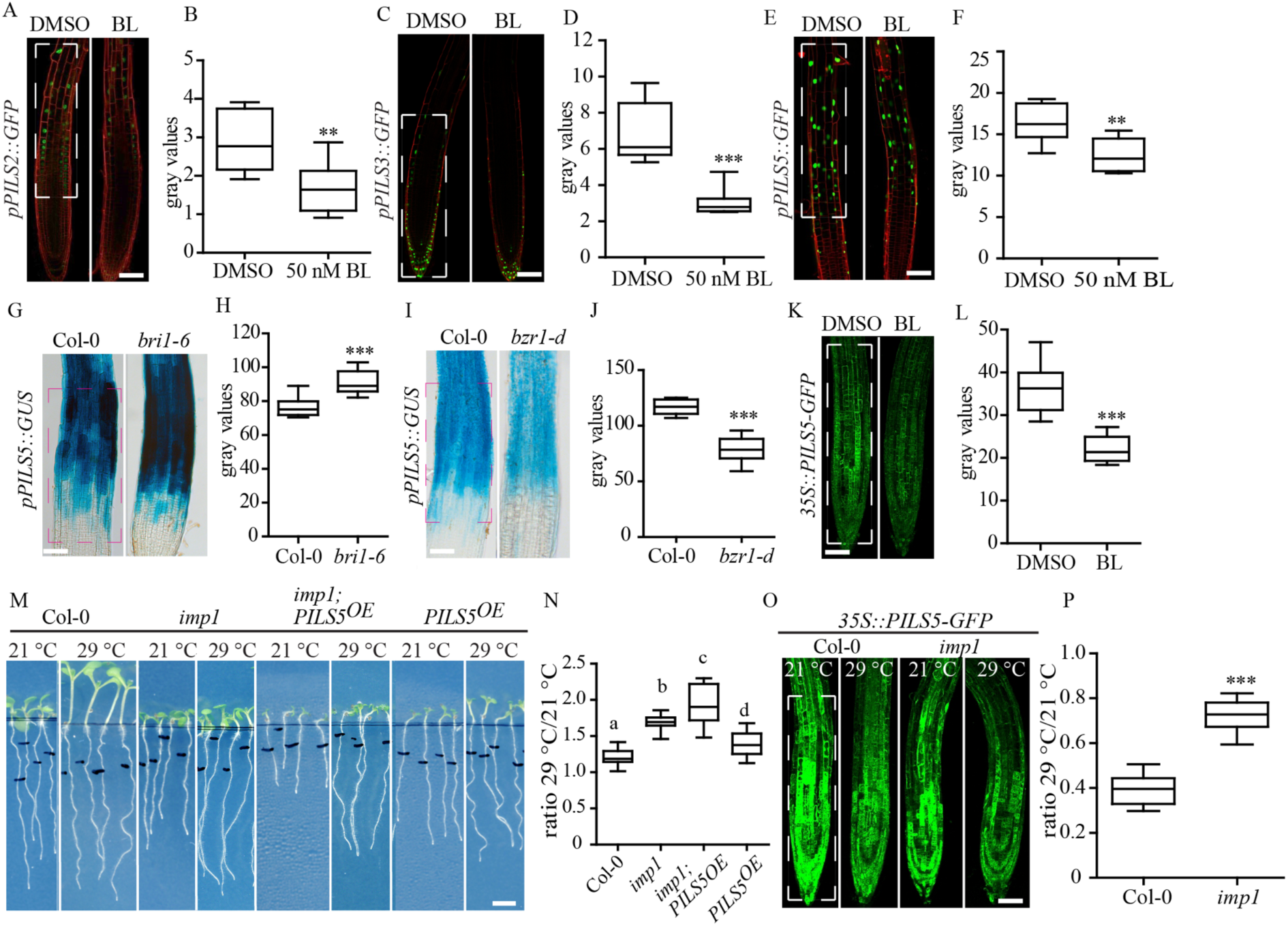
BR signaling represses PILS transcription and protein abundance. A-F, images and quantifications of *pPILS2::GFP* (A, B), *pPILS3::GFP* (C, D), and *pPILS5::GFP* (E, F) expression patterns in roots treated with DMSO or 50 nM BL for 12 h. Scale bar, 25 µm. (n = 8). G-J, GUS images and measurements of *PILS5* promoter expression in main root of *Col-0, bri1-6* (G, H) and *bzr1-d* (I, J). Scale bars, 140 mm. K-L, confocal images and quantification of PILS5-GFP fluorescence after treatment with DMSO or BL for 5 h. M and N, scanned images and quantifications (ratio) of the root segment grown after 3 d exposure to 21 °C (control) or 29 °C (HT). Scale bar, 30 mm. (n > 20). O and P, confocal images and quantifications of PILS5-GFP fluorescence in wild type and in *bri1* mutant after exposure to 21 °C (control) or 29 °C (HT) for 3 h. Scale bar, 25 µm. (n = 8). h: hours, d: days. Stars and letters indicate values with statistically significant differences (**P < 0.01, *** P < 0.001, student’s *t*-test (B, D, F, H, J, and P); P < 0.01, one-way ANOVA (N)). The dashed boxes represent the ROIs used to quantify signal intensity.

Next, we assessed whether BR signaling may also regulate PILS protein turnover, using lines that constitutively overexpress PILS-GFP proteins, under the control of the CaMV 35S promoter. PILS proteins, such as PILS2, PILS3, PILS5, and PILS6 were downregulated within hours of BL application (Figure 3K, 3L; Supplemental Figure 3E, 3F). This set of data suggests that BR signaling interferes with PILS function in a transcriptional and posttranslational manner.

The dual effect of BR signaling on *PILS* transcription and PILS protein abundance is reminiscent to the impact of high temperature, which also represses PILSes in a transcriptional and posttranslational manner (Feraru et al., 2019). High temperature-induced downregulation of PILS abundance elevates nuclear auxin input and increases root organ length (Feraru et al. 2019). Notably, BRI1-dependent BR signaling is also implied in root growth promotion under elevated ambient temperature (Martins et al., 2017). These independent findings prompted us to investigate whether BR signalling and PILS proteins jointly contribute to high temperature-induced root growth. Both *bri1* mutant and PILS overexpressing line display shorter roots compared to wild type under standard (21 °C) conditions. Even though absolute root length of *bri1* mutants and *PILS5*^*OE*^ lines remained also shorter when shifted to high temperature (29 °C) conditions, both lines showed a relative enhancement of root responses to high temperature when compared to wild type (Martins et al., 2017; Feraru et al., 2019; Figure 3M, 3N). The lines overexpressing PILS5 in *bri1* mutant background showed a similar relative enhancement of root response to high temperature (Figure 3M, 3N) as *bri1-6* and *PILS5*^*OE*^, proposing that BRI1 and PILS proteins are jointly implied in high temperature responses in roots. Hence, we next tested whether BR perception modulates PILS abundance under high temperature. We germinated *PILS5*^*OE*^ seedlings at 21 °C for 5 days and subsequently shifted the seedlings to 29 °C for three hours. As expected, the PILS5-GFP signal intensity strongly decreased in response to high temperature (Feraru et al., 2019; Figure 3O-3P). In contrast, genetic interference with *BRI1* impaired the high temperature-induced reduction of PILS5-GFP (Figure 3O-3P). This set of data indicates that BR signaling is implied in temperature-induced repression of PILS proteins.

### BR signaling modulates auxin signaling in a PILS-dependent manner

BR signaling impacts on ARF transcription factors (Vert et al., 2008; Sun et al., 2010; Cho et al., 2014; Oh et al., 2014) and induces auxin signaling in roots (Mouchel et al., 2006). Importantly, we noted that BL application does not only increase the auxin signaling output as previously shown (Mouchel et al., 2006; Figure 4K, 4M), but also decreased the fluorescence intensity of the nuclear auxin input marker DII-VENUS (Brunoud et al., 2012) within 1.5 to 3 hours (Figure 4A, 4B, 4F, 4G). This finding indicates that BL application increases the nuclear abundance of auxin and reveals an alternative, previously unanticipated BR-auxin crosstalk mechanism.

**Figure 4.**
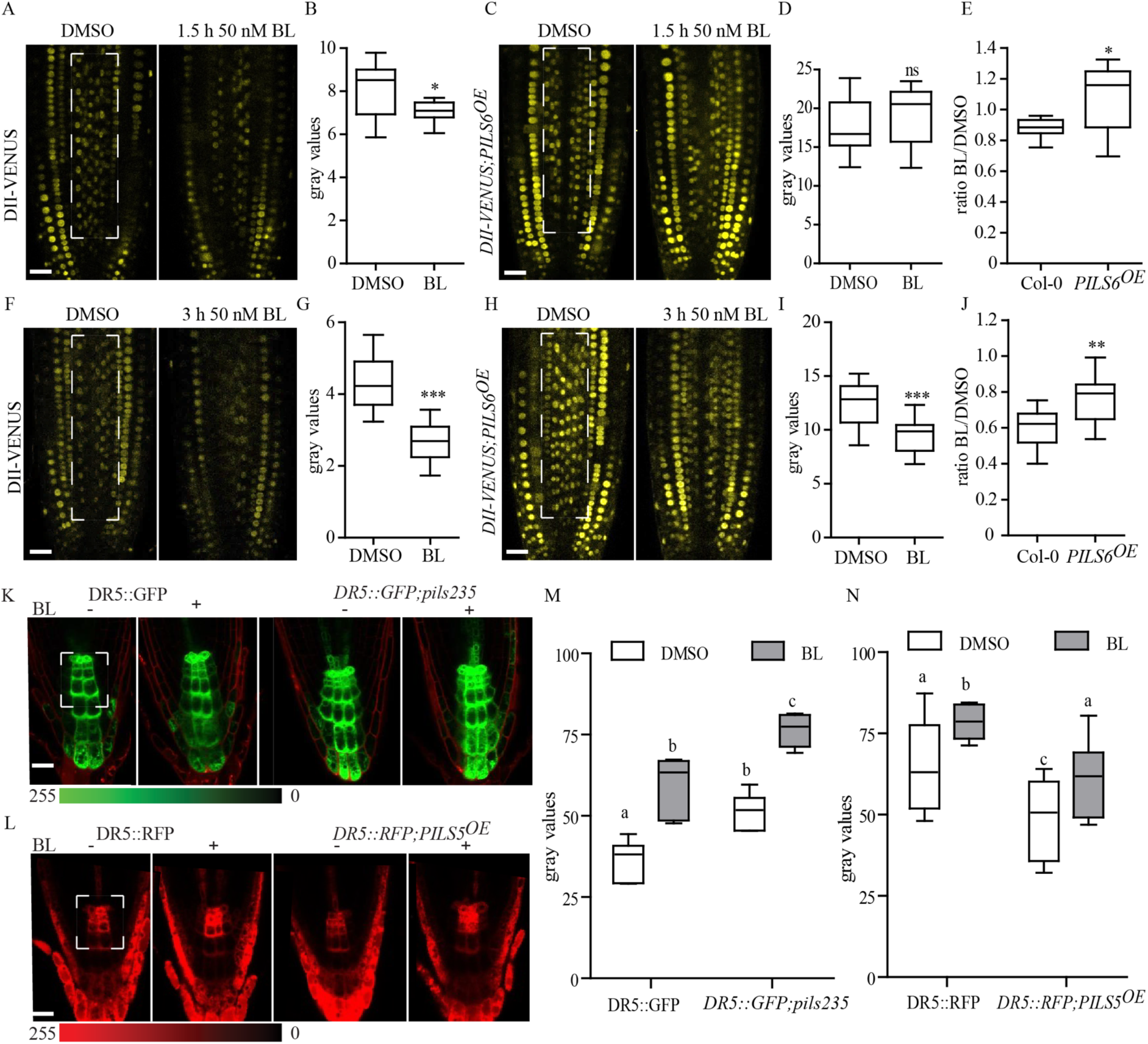
BR defines PILS-dependent nuclear abundance and signaling of auxin. A-J, confocal images (A, C, F, H) and absolute (B, D, G, I) or relative (E, J) quantifications of DII-VENUS in *Col-0* and in *PILS6*^*OE*^ treated with DMSO or 50 nM BL for 1.5 h (A-E) and 3 h (F-J). K-N, confocal images (K, L) and quantification (M, N) of *DR5::GFP* in *pils2 pils3 pils5* (*pils235)* (K, M) and *DR5::RFP* in *PILS5*^*OE*^ (L, N) roots exposed to DMSO or 50 nM BL. Scale bars, 25 µm. (n > 8). Stars and letters indicate values with statistically significant differences (*P < 0.05 **P<0.01 *** P < 0.001, student’s *t*-test (B, D, E, G, I and J), ns: no significant difference; P < 0.05, two-way ANOVA (M and N)). The dashed boxes represent the ROIs used to quantify signal intensity.

ER-localised PILS proteins repress nuclear availability and signaling of auxin (Barbez et al., 2012; Barbez et al., 2013; Beziat et al., 2017; Feraru et al., 2019), which prompted us to assess next whether BR-induced depletion of PILS proteins defines nuclear abundance of auxin. The BL-induced reduction of PILS6 protein abundance was relatively weak compared to the reduction of PILS3 and PILS5 proteins (Figure 3K, 3L; Supplementary Figure 3E, 3F). Hence, we tested if constitutive expression of PILS6 could partially counteract BR-dependent control of nuclear availability of auxin. The impact of BL on nuclear auxin input marker DII-VENUS was indeed reduced in *35S:PILS6-GFP* (*PILS6*^*OE*^) line when compared to wild type background (Figure 4A-4J). This set of data suggests that BR-dependent repression of PILS proteins modulates nuclear abundance of auxin.

The mutated mDII-VENUS is the auxin-insensitive version of DII-VENUS markers (Brunoud et al., 2012; Tan et al., 2007), disrupting the interaction between the DII domain, auxin, and the auxin receptors TIR1/AFBs. Prolonged (3 hours), but not short term (1.5 hours), exposure to BL induced a partial reduction in mDII-VENUS (Supplemental Figure 4A-4C). This unexpected sensitivity reminds of the high temperature effect, which also led to strong downregulation of DII-VENUS and comparably weaker depletion of mDII-VENUS (Feraru et al., 2019). Previous studies have suggested that mDII is insensitive to auxin (Brunoud et al., 2012; Liao et al., 2015), but under our conditions mDII-VENUS still remained partially sensitive to BR-(Supplemental Figure 4A-4C) or temperature-induced (Feraru et al., 2019) upregulation of nuclear auxin.

Next, we tested if the BR-reliant control of PILS-dependent nuclear abundance modulates auxin output signaling, by using the auxin responsive promoter DR5 transcriptionally fused to GFP (*DR5::GFP;* Benkova et al., 2003). While the sensitivity of *pils2, pils3, pils5* single or double mutant combinations were largely not distinguishable from wild type, we revealed that the BR-induced auxin signaling was markedly accelerated in *pils2-1 pils3-1 pils5-2* triple mutant roots (Figure 4K, 4M). This data suggests that PILS proteins redundantly contribute to BR responses.

As expected, BR-induced repression of PILS5 proteins (Figure 3K, 3L) also correlated with strongly increased nuclear auxin signaling (Figure 4L, 4N). Notably, absolute levels of *DR5::RFP* (Marin et al., 2010) remained quantitatively lower in the *PILS5*^*OE*^ when compared to the respective wild type seedlings (Figure 4N). This set of data suggests that BR-dependent repression of PILS5 modulates the nuclear availability and signaling of auxin.

### BR signaling modulates PILS-dependent organ growth rates

Our data proposes that BR signaling represses *PILS* expression and PILS protein abundance, which consequently increases the nuclear availability and signaling of auxin. To assess the potential developmental importance of this mechanism, we tested whether PILS proteins could define the sensitivity of root organ growth to BL application. BL treatment causes root growth reduction in wild type (Figure 5A, 5B). While the BL-sensitivity of *pils2, pils3, pils5* single or double mutant combinations were largely indistinguishable from wild type, we found that the total root growth of *pils2 pils3 pils5* triple mutant was hypersensitive to exogenous BL application (Figure 5A, 5B). In contrast, the constitutive expression of PILS5 induced hyposensitive root growth on BL (Figure 5C, 5D). BR signaling controls the cell cycle and subsequently root meristem size (Gonzales-Garcia et al., 2011). In agreement with the root length measurements, the negative impact of BL on meristem size was markedly amplified in *pils2 pils3 pils5* triple mutant and partially restored by constitutive PILS5 overexpression when compared wildtype meristems (Figure 5E-5G). These findings suggest that the BR-dependent control of PILS abundance contributes to root organ growth regulation. Similar to roots, dark grown hypocotyls of *pils2 pils3 pils5* triple mutant and *PILS5*^*OE*^ showed hyper-and hypo-sensitive growth responses to exogenously applied BL, respectively (Supplemental Figure 5A, 5B). Accordingly, this set of data proposes that PILS proteins are important integrators of phytohormonal crosstalk, allowing BR signaling to modulate nuclear abundance of auxin and organ growth rates.

**Figure 5.**
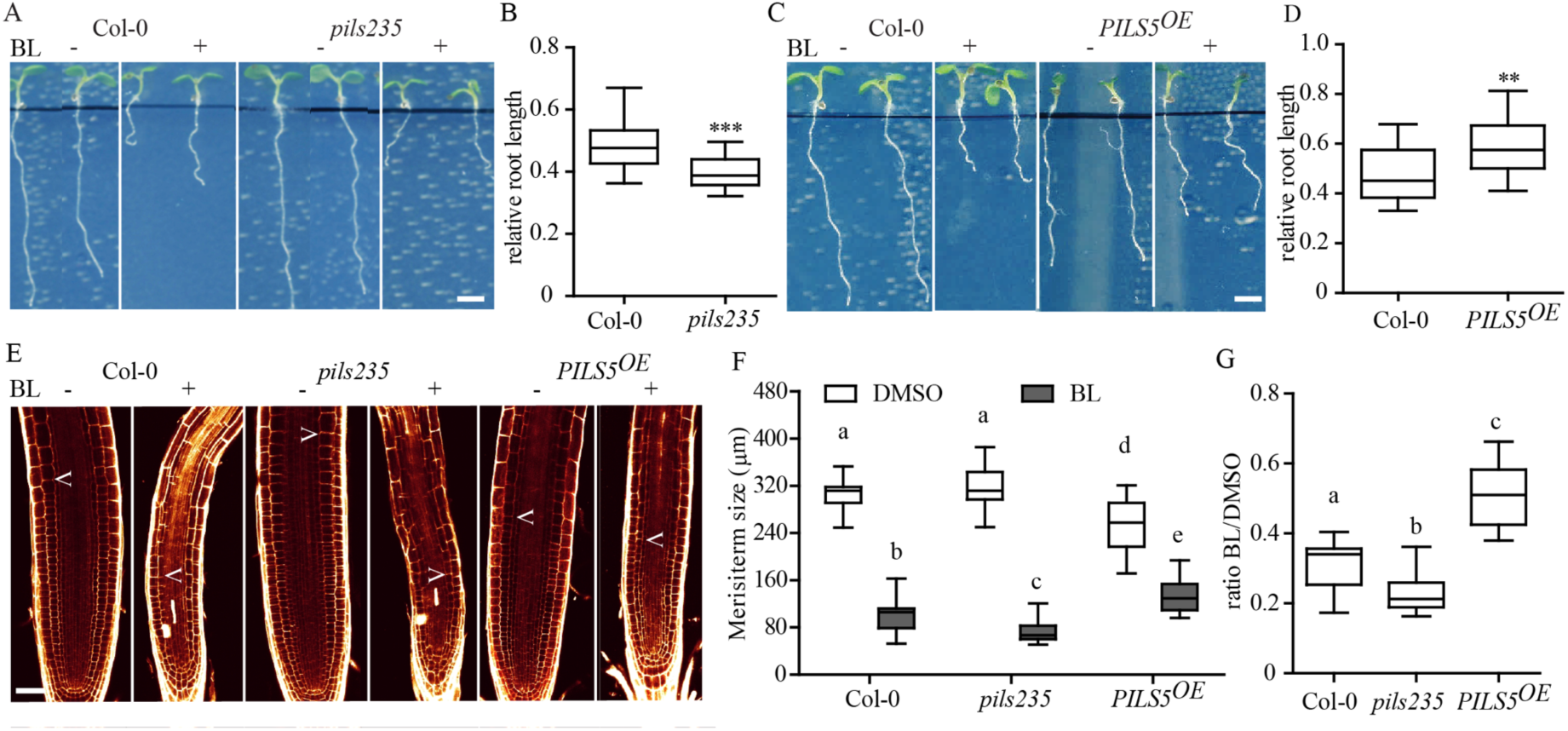
BR signaling modulates PILS-dependent root growth. A-D, images and quantifications of 6-d-old light-grown seedlings of *Col-0, pils235* (A and B), and *PILS5*^*OE*^ (C and D) germinated on plates with DMSO or 50 nM BL. Scale bar, 30 mm. (n > 30). E-G, confocal images and absolute (F) as well as relative (G) quantification of primary root meristem length of 6-d-old light-grown seedlings germinated on plates with DMSO or 50 nM BL. Scale bars, 25 µm. (n > 8). Stars and letters indicate values with statistically significant differences (**P < 0.01 ***P < 0.001, student *t*-test (B and D); P < 0.05, two-way ANOVA (F); one-way ANOVA (G)).

### A dynamic model on PILS-dependent crosstalk of auxin and brassinosteroid signaling

Auxin induces the expression of Aux/IAA genes that are potent transcriptional repressors of auxin signaling (Sauer et al., 2013), whereas BR promotes a negative effect on its own production (Wang et al., 2002). Accordingly, both auxin and BR demonstrate temporally delayed negative feedback on their activities, which could produce oscillations in both signaling circuits (Middleton et al, 2010; Allen and Ptashnyk, 2017). To illustrate dynamics of coupled systems, such as the auxin-BR crosstalk, we developed a computational model that implements these delayed negative feedback mechanisms on BR and auxin signaling (Fig 6A). Our model incorporates BRZ and ARF homodimers as well as ARF-BZR heterodimer moieties. In the first scenario the PILS-derived feedback is not considered and, thus, coupling between the two signaling circuits is solely through ARF-BZR heterodimers (Figure 6A). Model simulations are indeed capable of generating oscillations, however, auxin and BR oscillators varied in phase and, thus, the integrated signal (ARF-BZR heterodimers) were not synchronized (Fig. 6A; Supplementary Fig 6A and 6E). Next, we incorporated PILS function in the model, taking into account the here revealed negative effect of BR on PILS protein stability and gene expression, as well as the positive impact of auxin on *PILS* transcription (Barbez et al., 2012). The model simulation predicted a tight phase-locking and consequently synchronization of two oscillators (Fig. 6B and Supplementary Fig. 6B and 6E). Furthermore, balanced levels of PILS seems critical to sustain the synchrony between BR-auxin signaling, since a gradual overexpression of PILS caused a phase drift in our model, reminiscent of decoupled oscillations (Supplementary Fig. 6C-6E).

**Figure 6.**
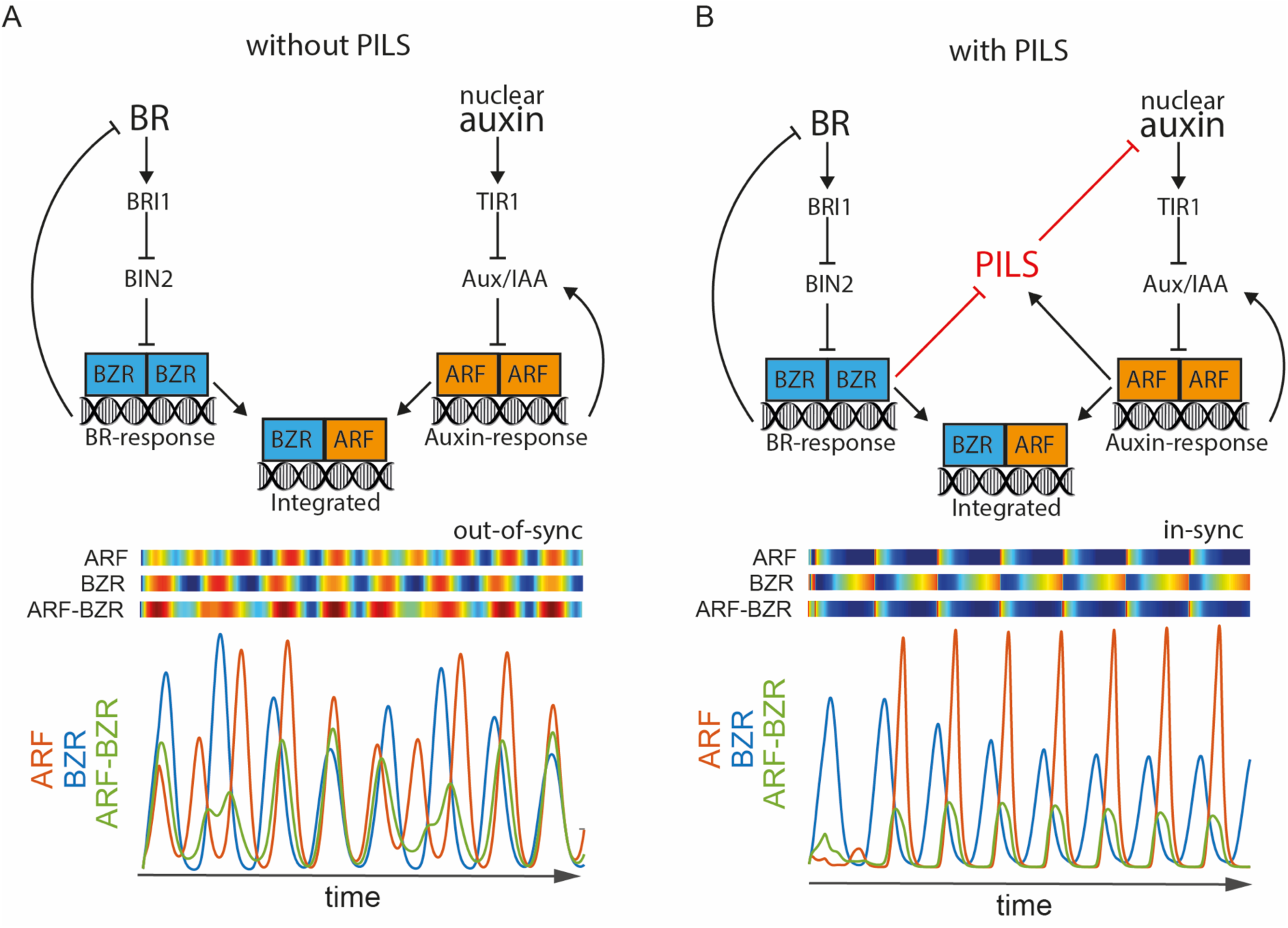
A computer model predicts PILS-dependent synchronization of BR and auxin responses. A and B, schematics of BR-auxin oscillatory mechanism without (A) and with PILS-dependent feedback (B), top panel. Computer model simulations are shown as heat maps (blue to red) and corresponding time-lapse curves with activity peaks for BZR (blue), ARF (red) and ARF-BZR (green) (bottom panel).

Taken together, our model suggests that BR and auxin signaling are coupled through PILS proteins, thereby acting in synchrony.

## Discussion

BRs and auxin play overlapping roles in plant growth and development and, intriguingly, many target genes of BR and auxin signaling are overlapping. Increased auxin levels saturate the BR-stimulated growth response and greatly reduce the BR effects on gene expression (Nemhauser et al., 2004). BR-dependent BIN2 signaling component and BZR1/2 transcription factors, have been previously shown to directly regulate the ARF transcription factors (Vert et al., 2008; Sun et al., 2010; Cho et al., 2014; Oh et al., 2014), which are key components in realizing the transcriptional output of auxin. Most intriguingly, BZR1 and ARF6 transcription factors directly interact (Oh et al., 2014) and this direct crosstalk mechanism is thought to integrate and specify BR and auxin signaling output. Here, we reveal a higher molecular complexity in the BR-auxin crosstalk, indicating that BR does not only modulate auxin output signaling, but also nuclear input of auxin.

We have previously shown that PILS proteins determine intracellular accumulation of auxin at the ER, decrease cellular sensitivity to auxin, and negatively impact on nuclear availability, as well as signaling of auxin (Barbez et al., 2012; Barbez et al., 2013; Beziat et al., 2017; Feraru et al., 2019). Mechanistically, we assume that PILS proteins retain auxin in the ER and thereby reduce the diffusion of auxin from the cytosol into the nucleus. Our forward genetic screen reveals that the BR signaling affects the PILS-reliant traits by restricting the abundance of PILS proteins and thereby increasing nuclear input and signaling rates of auxin. We, accordingly, revealed an alternative, unanticipated BR-auxin crosstalk mechanism, which may also explain how BR sensitizes seedlings to auxin (Vert et al., 2008).

Auxin signaling itself stimulates *PILS* gene expression (Barbez et al., 2012), presumably acting as a negative feedback mechanism on auxin signaling. Additionally, external cues, such as light, modulate *PILS* transcription in a Phytochrome Interacting Factors (PIFs)-dependent manner and, thereby, define differential growth responses in apical hooks (Beziat et al., 2017). Here, we show that BR signaling represses expression of *PILS2, PILS3* and *PILS5* genes. We also show that *BZR1* overexpression limits *PILS5* promoter activity in roots, which agrees with *PILS2* and *PILS5* being listed as direct targets of BZR transcription factors (He et al., 2005; Chaiwanon and Wang, 2015).

Besides the effect on *PILS* transcription, we show that BR signaling posttranslationally restricts PILS protein abundance. This finding is reminiscent to the effect of increased ambient temperature, which also transcriptionally and posttranslationally limits PILS protein abundance and, thereby, increases nuclear abundance and signaling of auxin (Feraru et al., 2019). Moreover, high temperature induces root organ growth in a BR-(Martins et al., 2017) and auxin-dependent manner (Feraru et al., 2019). Here we illustrate that high temperature-induced repression of PILS5 protein requires BR signaling. Accordingly, we propose that the here described phytohormonal crosstalk mechanism has developmental importance, integrating BR signaling with PILS-dependent auxin responses and root organ growth.

BR does not regulate the expression of PIN intercellular transport components (Hacham et al., 2011) and its effect on root meristem size has been proposed to be independent of auxin (reviewed in Tian et al., 2018). On the other hand, the balance between BR and auxin levels is known to be required for optimal root growth (Chaiwanon and Wang, 2015). Here, we show that PILS proteins are involved in a transverse BR-auxin crosstalk mechanism (Supplemental Figure 6), which quantitatively contributes to meristematic activity and overall root growth rates. Untreated *pils2 pils3 pils5* triple mutant and *PILS5*^*OE*^ tendentially display bigger and smaller meristems when compared to wild type, respectively. Most strikingly, the application of BR reverses the meristem regulation, leading to smaller and bigger root meristems in *pils2 pils3 pils5* triple mutant and *PILS5*^*OE*^ lines, respectively. A similar trend was observed for high temperature-induced root organ growth (Feraru et al., 2019), which involves the here identified BR-auxin crosstalk. Accordingly, we assume that the BR effect on PILS proteins not only quantitatively set auxin signaling rates, but also qualitatively define the hormonal crosstalk between BR and auxin. This assumption is also in agreement with our computational model, predicting that PILS proteins coordinate BR and auxin signaling. We, hence, anticipate that BR-dependent control of PILS activity may have additional wide-spread importance during plant growth and development by synchronizing BR and auxin signaling responses.

## Material and Methods

### Plant material and growth conditions

*Arabidopsis thaliana* ecotype *Columbia-0* (*Col-0*) and *Landsberg erecta* (*Ler*) were used for experiments. The following lines have been described previously: *bri1-5* and *bri1-6* (*Enkheim-2*) (Noguchi et al; 1999), *pBRI1::BRI1-GFP* (Friedrichsen et al; 2000), *bzr1-d* (Wang, et al 2002), *pDR5rev::GFP* (Friml et al; 2003), *pDR5rev::mRFP1er* (Marin et al; 2010), *p35S::GFP-PILS2* (*PILS2*^*OE*^), *p35S::GFP-PILS3* (*PILS3*^*OE*^), *p35S::PILS5-GFP* (*PILS5*^*OE*^), *p35S::PILS6-GFP* (*PILS6*^*OE*^), and *p35S::PILS5-GFP;pDR5rev::mRFP1er, pils2-1, pils5-2* (Barbez et al., 2012), *pPILS2, 3, and 5::GFP/GUS-NLS, pils3-1* (Béziat et al, 2017a), *DII-VENUS* and *mDII-VENUS* (Brunoud et al., 2012), *DII-VENUS;PILS6*^*OE*^ *and mDII-VENUS;PILS6*^*OE*^ (Feraru et al., 2019). Multiple mutants and marker lines were generated by crossing.

Seeds were stratified at 4°C for 2 days in dark. Seedlings were grown vertically in Petri dishes on ½ Murashige and Skoog (MS) in vitro plates supplemented with 1% sucrose and 1% agar (PH5.9). Plants were grown under the long-day (16 h light/8 h dark) condition at 21 (±1) °C. For treatments, 5-or 6-d-old seedlings were incubated for 5 h or 12 h in solid and /or liquid MS medium containing the indicated concentrations of 24-Epibrassinolide (BL) (Invitrogen; 1 or 10 mM in DMSO solvent) or germinated for five or six days on MS medium supplemented with BL at 100 nM and 50 nM, respectively. For high temperature (HT)-related experiments, two growth cabinets were equipped with overhead LED cultivation lights (Ikea, 703.231.10), at an irradiance of 150 µmol/m^−2^s^−1^, and set at 21 °C (control) or 29 °C (HT treatment) under long-day conditions. For microscopy, the seedlings were grown on vertically oriented plates for five days under 21 °C, and then kept under 21 °C (control) or transferred to 29 °C (HT) for 3 h. For root growth analysis, seedlings were grown for seven days under 21 °C (control) and for four days under 21 °C followed by three days under 29 °C (HT).

### Forward genetic screen and mapping

To identify modulators of PILS5, *35S::PILS5-GFP (PILS5*^*OE*^) seedlings descended from 3000 ethyl methanesulfonate (EMS) (0.3%) mutagenized M1 plants were analyzed for the dark-grown hypocotyl phenotype. The *imp1* mutant was mapped on the upper arm of chromosome 4 between nga1107 (18.096 Mb) and T5J17-16 (18.570 Mb). A total number of 87 recombinants from the F2 cross between *imp1* (*Columbia* background) and *Landsberg erecta* were used. For *Columbia*/*Landsberg erecta* polymorphism information, the Monsanto Arabidopsis Polymorphism and the *Ler* Sequence Collection (Cereon Genomics) were used. For information regarding single nucleotide polymorphisms and insertions/deletions, the Arabidopsis Information Resource (http://www.Arabidopsis.org) was used.

### Next generation sequencing

The genomic DNA of *imp1* was prepared for next generation sequencing. Fifteen individuals of F2 progeny derived from cross of *imp1* with the *Col-0* were selected based on the dark-grown hypocotyl phenotype. The selected seedlings were transferred to soil. Subsequently, leaf tissue from 3-w-old plants was harvested for DNA isolation. Genomic DNA extraction was performed using DNeasy plant mini kit (QIAGEN) according to the manufacturer’s handbook. The DNA samples were sent to BGI Tech (https://www.bgi.com) for Whole Genome Re-sequencing using Illumina’s HiSeq 2000.

### Phenotype analysis

For hypocotyl analysis, seeds on plates were exposed to light for 8 h at 21 °C, cultivated in the dark at 20 °C, and scanned at 4-or 5-d-old. For analysis of root length, 6-d-old seedlings on solvent or treatment containing plates were scanned. For root response to HT, 4-d-old root tips of seedlings grown under 21 °C were marked before the transfer for three additional days under 21 °C (control) or 29 °C (HT). Only the root segment grown after the transfer was measured. Plates were scanned with an Epson Perfection V700 scanner. Hypocotyl and root lengths were measured with the ImageJ (http://rsb.info.nih.gov/ij/) software.

### Quantification of root meristem

Root meristems of 6-d-old seedlings grown on solid plates with DMSO or 50 nM BL were imaged with a Leica TCS SP5 confocal microscope. Seedlings were stained with Propidium Iodide (0.02mg/mL) before imaging. The meristem size was defined as the distance between the quiescent center and the first rectangular cortical cell (Löfke et al., 2015). Leica software (LAS AF Lite) was used for quantification.

### GUS staining

GUS staining was performed and quantified as described previously (Beziat et al., 2017b). The whole seedlings of 5-d-old dark grown or 6-d-old light grown with or without BL treatment were harvested to wells containing 1 ml of cold 90% acetone and incubated for 30 minutes on ice. The rehydrated seedlings were mounted in chloralhydrate for analysis by light microscopy (Leica DM 5500) equipped with a DFC 300 FX camera (Leica). To quantify the signal intensity, a Region Of Interest (ROI) was defined to capture the most representative signal distribution. This region is indicated in the figures and was kept constant (size and shape) for all analyzed samples.

### Confocal Microscopy

5-or 6-d-old *35S::PILS5-GFP* and *pDR5::GFP/RFP* seedlings in *Col-0* or mutant backgrounds were imaged with a Leica SP5 or Leica SP8 (Leica). Fluorescence signals for GFP (excitation 488 nm, emission peak 509 nm), mRFP1 (excitation 561 nm, emission peak 607 nm) and propidium iodide (PI) staining (excitation 536 nm, emission peak 617 nm) were detected with 20× (water immersion) or 63× (water immersion) objective. To image *DII-VENUS* and *mDII-VENUS*, Leica TCS SP8 equipped with a white laser was used, allowing us to separate GFP and YFP fluorophores. The fluorescence signal intensity (mean gray value) of the presented markers was quantified using the Leica software.

### Statistical analysis

Means and standard errors were calculated and the statistical significance was evaluated using the Graph Pad Prism5 (http://www.graphpad.com) software. The significance of the data was evaluated using the student *t*-test in the case of two columns comparisons. One-way ANOVA followed by Tukey’s test was performed in the case of the multiple columns comparisons procedure. Two-way ANOVA followed by Bonferroni post-tests was carried out to compare two different genotypes at different treatments.

Representative data are shown throughout the text. All experiments have been performed in at least three replications.

### Computational methods

The dynamics of all components built in the model was simulated using delayed differential equations (DDEs) implemented in Matlab Inc. The Matlab-derived dde23 solver (https://www.mathworks.com/help/matlab/ref/dde23.html) was used to obtain direct solutions of DDEs. All simulations were performed until 300 steps to account for multiple oscillations. Overall, our model incorporates BR and auxin signalling pathways (Nemhauser, J. L., et al, 2004; Tian H, et al, 2018) with an addition of the here revealed BR-dependent regulation of PILS on transcriptional and post-translational levels.

#### Brassinosteroid signalling branch modelling

BR synthesis is inhibited by by its own signalling, which is implemented by BZR dimers (BZR^D^) with delay *τ*:

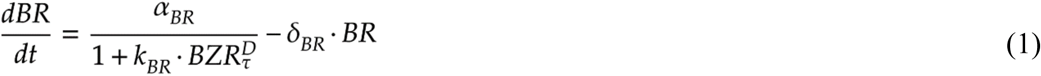

where *α*_*BR*_ are BR production rate and *k*_*BR*_ is rate of repression mediated by BZR^D^ and *δ*_BR_ is a BR turnover rate.

BR perception is known to define BZR activity by inhibiting BIN2 phosphorylation (Cho, H., et al, 2014). Hence, we included the BIN2 regulation into our model by:

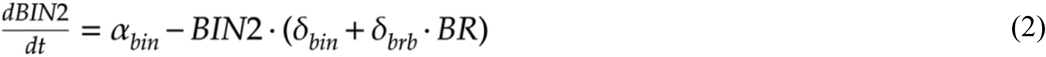

where *α*_*bin*_ and *δ*_bin_ are BIN2 production and degradation rates and *δ*_brb_ denotes the rate of BR-dependent BIN2 de-phosphorylation. Next, BIN2 interferes with nuclear BZR activity and thus negatively affects levels of BZR^D^:

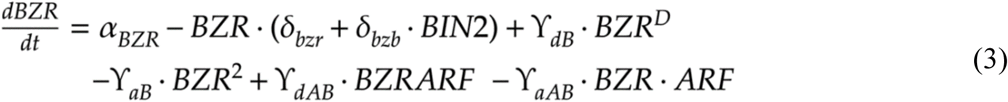

where *α*_*BZR*_ and *δ*_bzr_ are BZR production and degradation rates and *δ*_bzb_ denotes the rate of BIN2-dependent repression of BZR. *γ*_*dB*_ and *γ*_*dAB*_ stand for dissociation rates of BZR^D^ and BZRARF^D^ dimers whereas *γ*_*aB*_ and *γ*_*aAB*_ are association rates of these dimers. Note that BR steers a delayed negative feedback on its own production eq. 1-3.

Furthermore, species of BZR^D^ and BZRARF^D^ are given by following formulas:

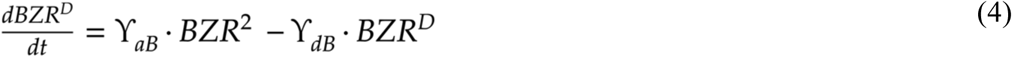

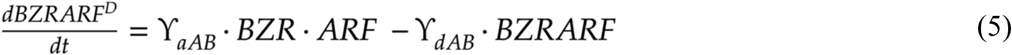

#### Auxin signalling branch modelling

Nuclear auxin (A) is restricted by PILS auxin transport facilitators (Barbez et al., 2012; Feraru et al., 2019):

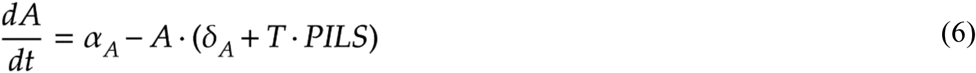

where *α*_*A*_ and *δ*_A_ are production and degradation constants of auxin and T is PILS transport coefficient. The dynamics of auxin signalling repressors (AUX/IAA) (Middleton AM, et al, 2010) are modelled by combining ARF-mediated transcription, translation and auxin-dependent degradation in the following formula:

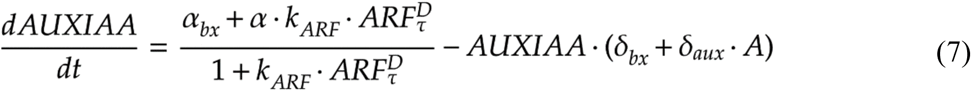

where *α*_*bx*_ and *δ*_bx_ are basal production and degradation constants of AUX/IAA (AUXIAA). *α* denotes the ARF-dependent transcription rate times amount of ARF homodimers (ARF^D^) and *δ*_aux_ is an auxin-dependent degradation rate. *k*_*ARF*_ is promoter association constant of ARF^D^ Next, ARF monomers (ARF) are described by the following mathematical equation:

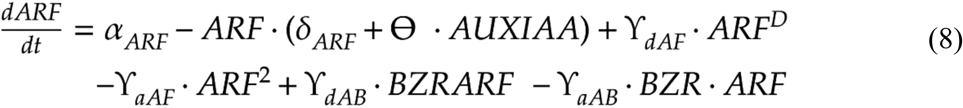

*α*_*ARF*_ and *δ*_ARF_ are basal production and degradation rates of ARF monomer and θ represents AUX/IAA-dependent ARF sequestering that leads to negative feedback on AUX/IAA levels. *γ*_*dAF*_ and *γ*_*aAF*_ stand for dissociation and association rates of ARF dimers (ARF^D^) that follow the formula:

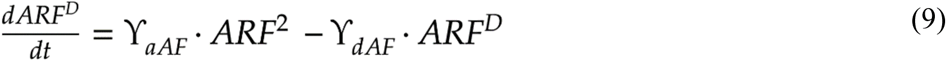

Finally, PILS protein levels are coupled to BR and auxin signalling pathway through transcription and degradation and follow this formula:

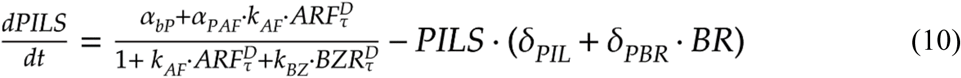

Where *α*_*bP*_ and *δ*_PIL_ are basal production and degradation rates of PILS proteins, respectively. *α*_*pAF*_ denotes the ARF-dependent transcription rate and *δ*_PBR_ is an BR-dependent degradation rate of PILS. *k*_*AF*_ is association constant of ARF^D^ to PILS promoter and *k*_*BZ*_ is a rate of repression mediated by BZR dimers.

Parameters of the computer model for all four conditions (without PILS, with PILS, PILS 5-fold OX and 100-fold PILS OX) are summarized in the following table:

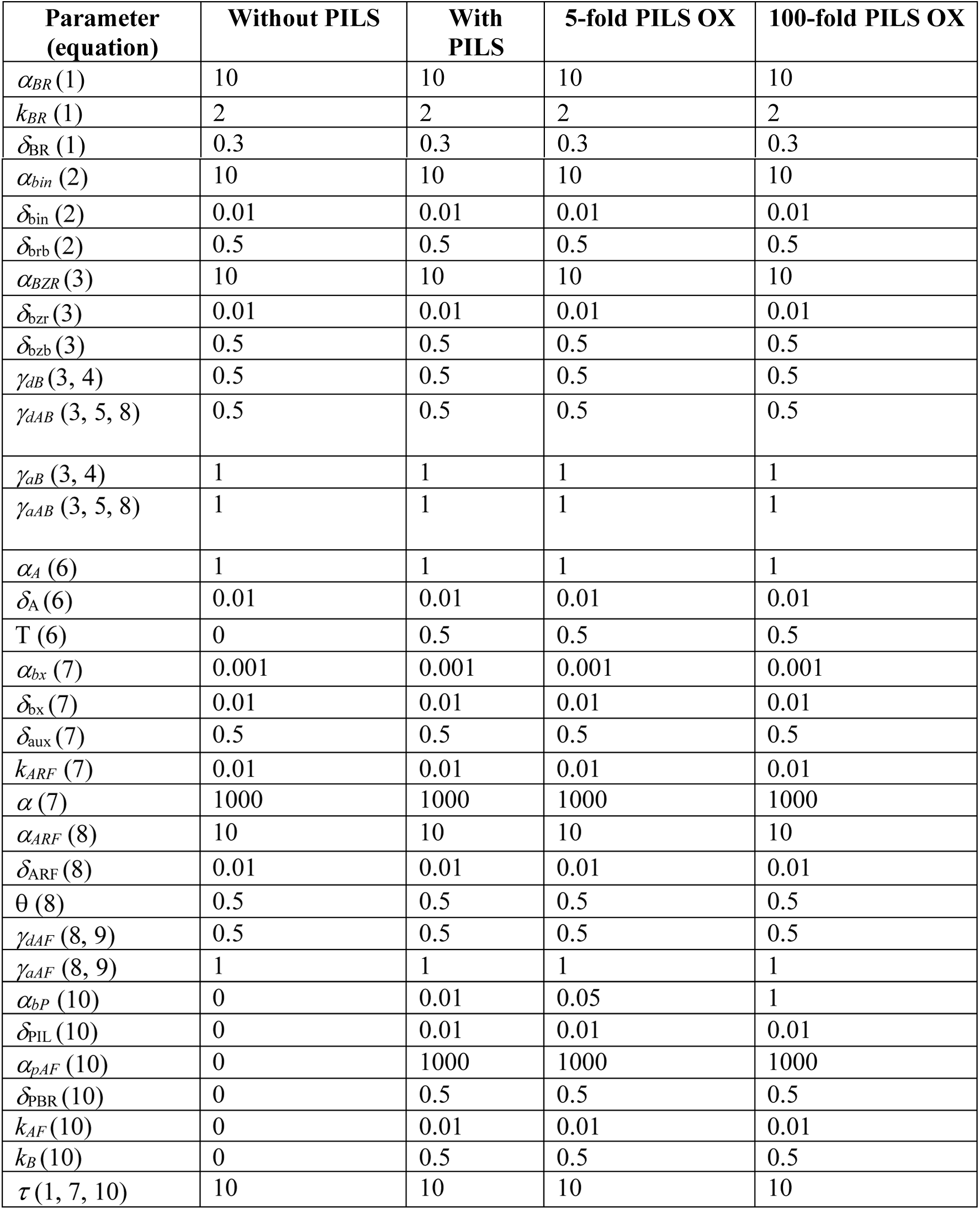

#### Phase difference calculations -synchrony measure

For each time-dependent solution of ARF^D^ and BZR^D^, amplitudes and periods were calculated, using **peakfind** function (Matlab Inc.) and subtracted to estimate phase differences between two oscillators. The phase differences were plotted, using violin plot function in Matlab (https://www.mathworks.com/matlabcentral/fileexchange/45134-violin-plot) together with probability density distributions performed with **histfit** function (https://www.mathworks.com/help/stats/histfit.html). The large variation in phases (broader distribution) indicates that two oscillatory pathways are out-of-sync, whereas sharper distributions reflects near-perfect synchrony between the two signalling pathways.

## Acknowledgements

We are grateful to J. Friml, Russinova E, R. Strasser, and T. Vernoux, for providing the published material; the BOKU-VIBT Imaging Center for access and M. Debreczeny for expertise; Elizabeth Sarkel and Elke Barbez for critical reading. This work was supported by the Vienna Science and Technology Fund (WWTF) (to J.K.-V.), Austrian Science Fund (FWF) (P26591-B16 to J.K.-V.), European Research Council (AuxinER -ERC starting grant 639478 to J.K.-V.), as well as FWF-Hertha Firnberg and Elise Richter (T728-B16 and V690-B25 to E.F.) and the China Scholarship Council (CSC) (predoctoral fellowship to L.S.), and Programa de Atracción de Talento 2017 (Comunidad de Madrid, 2017-T1/BIO-5654 to K.W.).

---

**Supplementary Figure 1.**
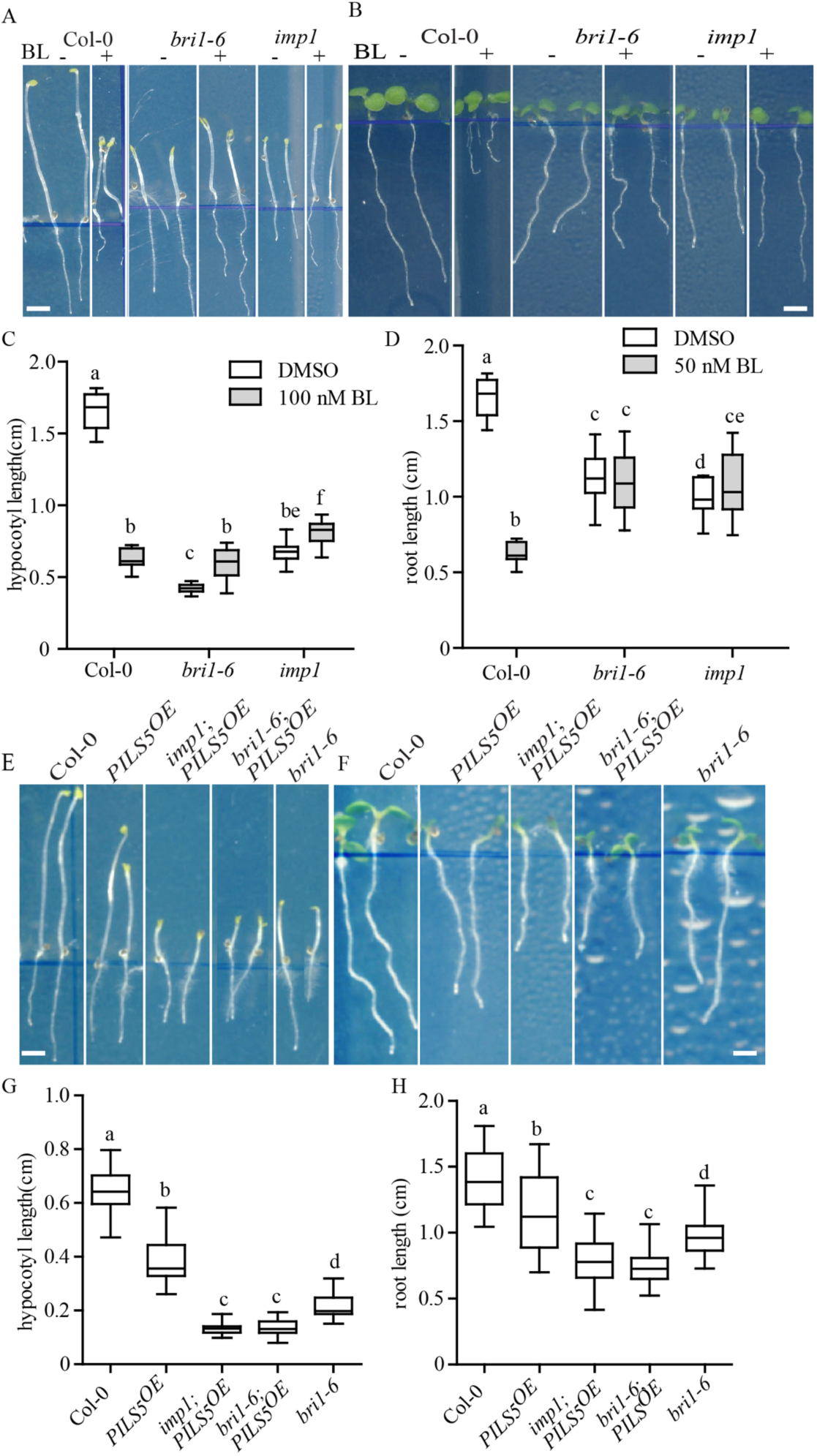
Impaired BR perception enhances *PILS5*^*OE*^ phenotypes. A-D, images and quantifications of 5-d-old dark-grown (A, C) and 6-d-old light-grown (B, D) seedlings of *Col-0, bri1-6* and *imp1* germinated on DMSO, 100 nM (A, C), or 50 nM BL (B, D). Scale bar, 30 mm. (n > 30). E-H, scanned images and quantifications of 5-d-old dark-grown (E, G) and 6-d-old light-grown (F, H) seedlings of wild type, *PILS5*^*OE*^, and *bri1* mutants. Scale bar, 30 mm. (n > 25). Letters indicate values with statistically significant differences (P < 0.05, two-way ANOVA (A-D); P < 0.01, one-way ANOVA (E-H)).

**Supplementary Figure 2.**
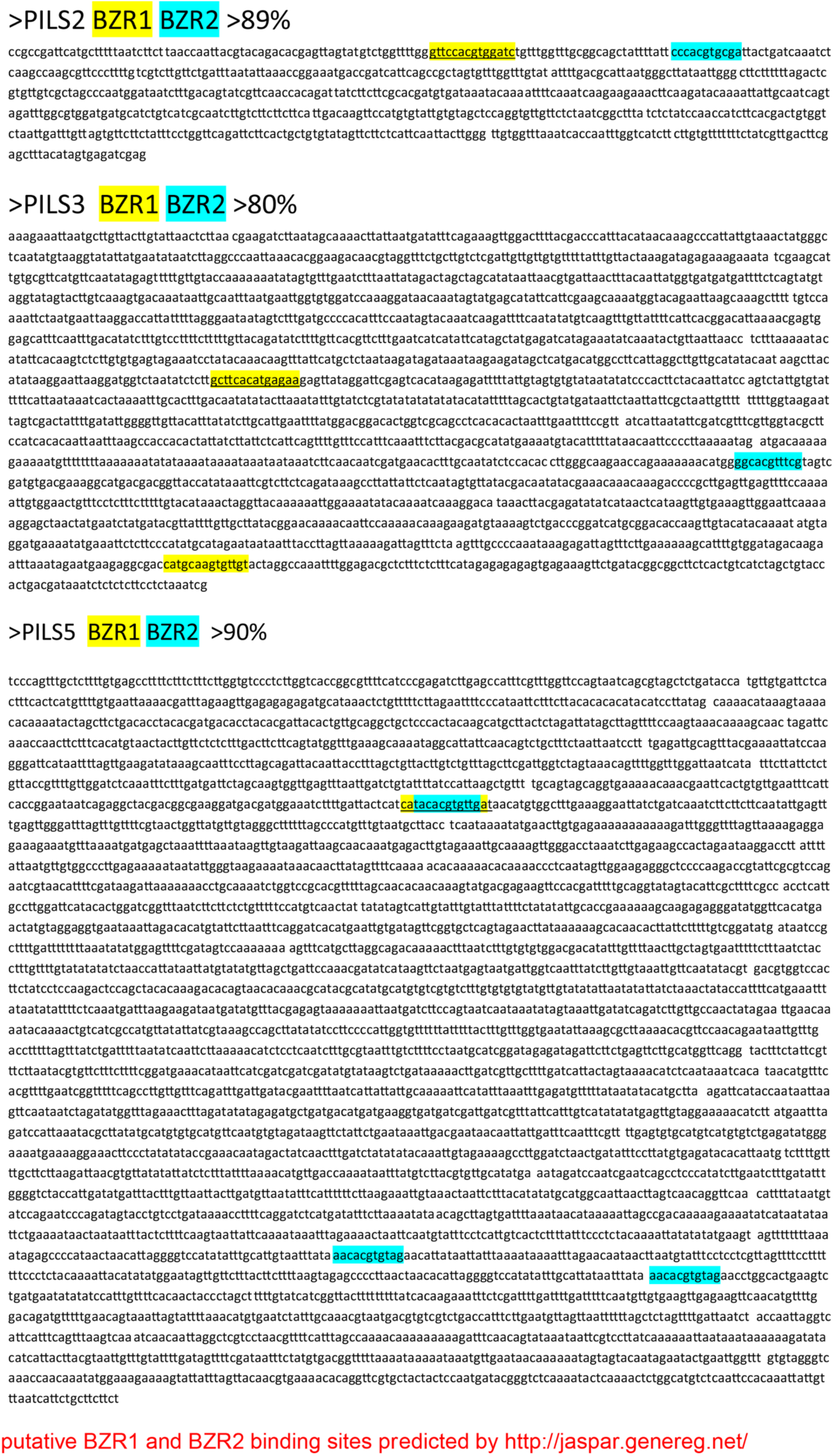
Predicted BZR1 and BZR2 binding motifs in the promoters of *PILS2, PILS3*, and *PILS5*. *in silico* predicted G boxes are depicted in yellow (for BZR1) and blue (for BZR2) in *PILS2, PILS3* and *PILS5* promoter sequences. The calculated probability of binding for *pPILS2* (p > 89%), *pPILS3* (p >80%), and *pPILS5* (p > 90%) is indicated.

**Supplementary Figure 3.**
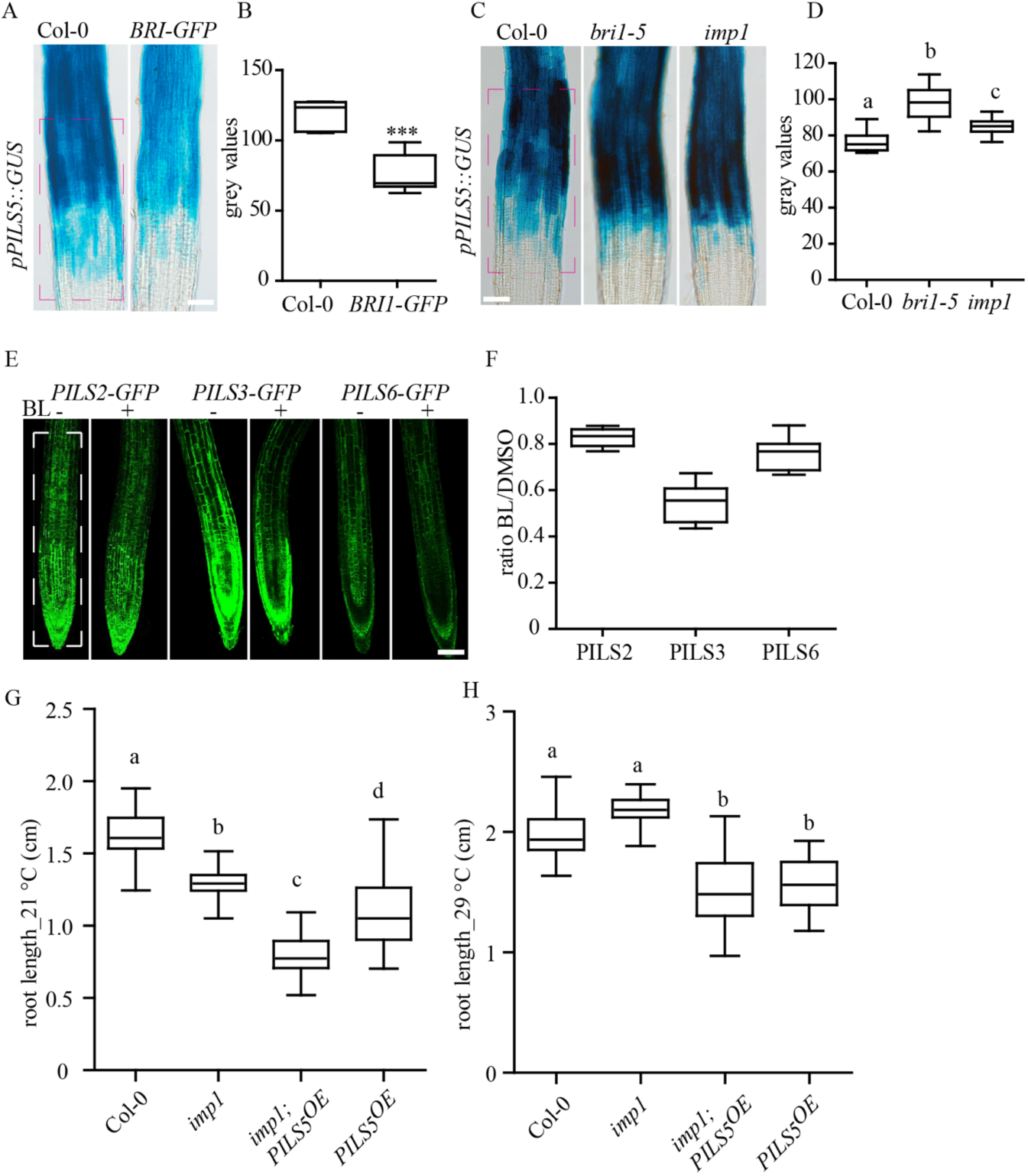
BR signalling negatively impacts on the PILS transcription and protein abundance. A-D, GUS images (A, C) and quantifications (B, D) of *pPILS5::GUS* expression pattern in roots of wild type and BRI1-GFP overexpressor (A, B), *bri1-5* and *imp1* (C, D). Scale bars, 140 mm. (n = 8). E and F, confocal images (E) and quantifications (F) of *35S::GFP-PILS2, 35S::GFP-PILS3*, and *35S::PILS6-GFP* treated with DMSO or 50 nM BL for 5 h. Scale bar, 25 µm. (n = 8). The dashed boxes represent the ROIs used to quantify signal intensity. Stars and letters indicate values with statistically significant differences (***P < 0.001, student *t*-test (B and D); P < 0.05, one-way ANOVA (F)).

**Supplementary Figure 4.**
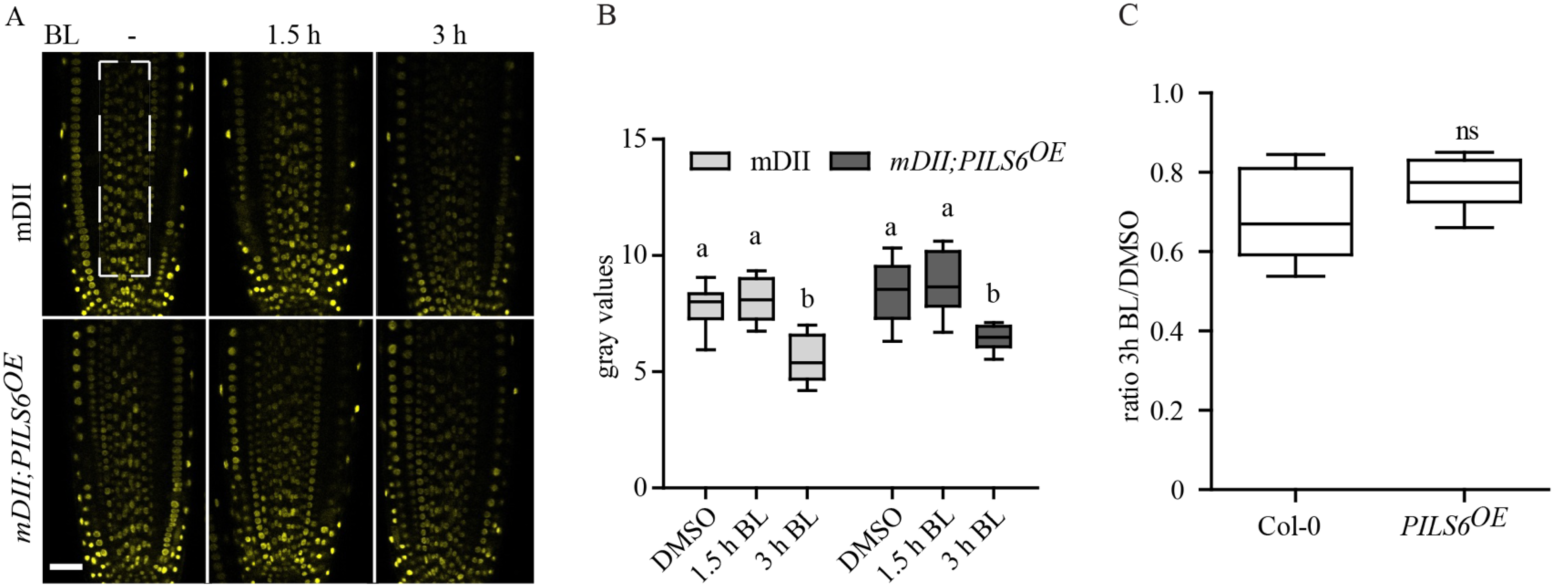
mDII is partially sensitive to BL. A-C, confocal images (A) and quantifications (B, C) of mDII-VENUS in wild type and in *PILS6*^*OE*^ after 1.5 h and 3 h of BL treatment. Scale bars, 25 µm. (n > 10). Letters indicate values with statistically significant differences (P < 0.001, two-way ANOVA (B); student’s *t*-test (C), ns: no significant difference). The dashed boxes represent the ROIs used to quantify signal intensity.

**Supplementary Figure 5.**
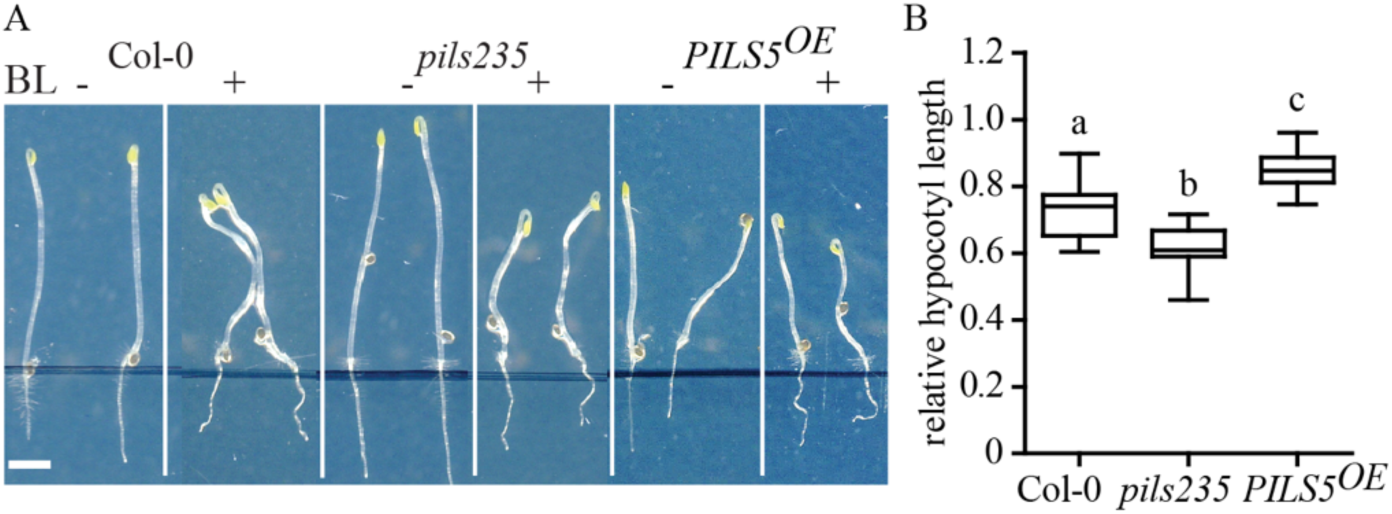
PILS proteins define BR-sensitive hypocotyl growth. A and B, scanned images (A) and quantifications (B) of 5-d-old dark-grown hypocotyls of *Col-0, pils2 pils3 pils5 (pils235)*, and *PILS5*^*OE*^ germinated on plates with DMSO or 100 nM BL. Scale bar, 30 mm. (n > 30). Letters indicate values with statistically significant differences (P < 0.01, one-way ANOVA).

**Supplementary Figure 6.**
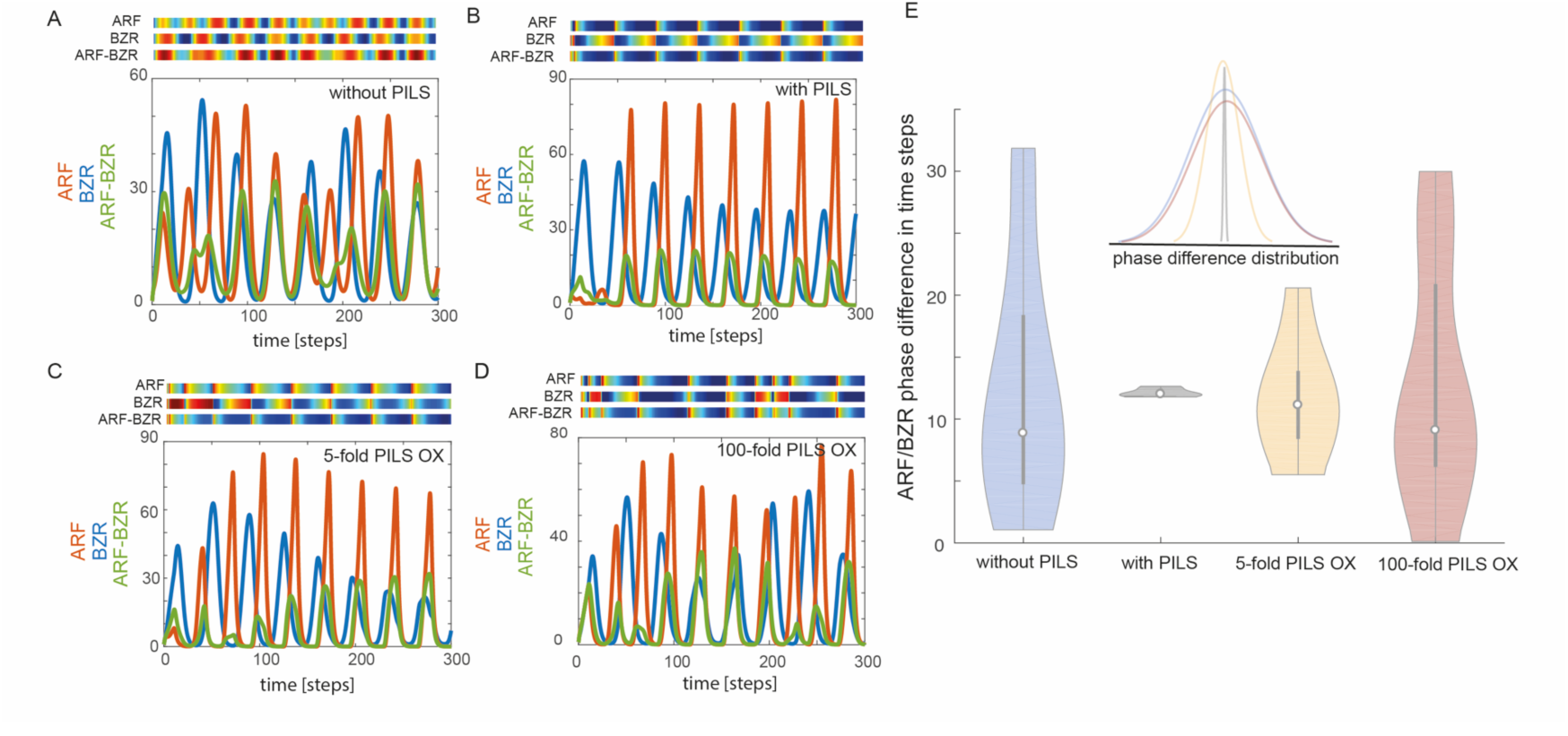
Simulations of computer model predicts the phase locking between BR and auxin signaling outputs through PILS dependent auxin transport. A-D, model simulations without PILS-(A), with PILS-dependent feedback (B) (corresponds to Figure 6A and B), as well as with mild (5-fold) PILS overexpression (C) and strong (100-fold) PILS overexpression (D). Graphs depict time-lapse for BZR (blue), ARF (red) and ARF-BZR (green) levels and corresponding heat maps (blue to red) for peak responses. E, Violin plots show phase difference between BZR and ARF oscillation, corresponding to model simulations (A-D).

